# A phenomics approach for *in vitro* antiviral drug discovery

**DOI:** 10.1101/2021.01.13.423947

**Authors:** Jonne Rietdijk, Marianna Tampere, Aleksandra Pettke, Polina Georgieva, Maris Lapins, Ulrika Warpman Berglund, Ola Spjuth, Marjo-Riitta Puumalainen, Jordi Carreras-Puigvert

## Abstract

**Background:** The current COVID-19 pandemic has highlighted the need for new and fast methods to identify novel or repurposed therapeutic drugs. Here we present a method for untargeted phenotypic drug screening of virus-infected cells, combining Cell Painting with antibody-based detection of viral infection in a single assay. We designed an image analysis pipeline for segmentation and classification of virus-infected and non-infected cells, followed by extraction of morphological properties.

**Results:** We show that the methodology can successfully capture virus-induced phenotypic signatures of MRC-5 human lung fibroblasts infected with Human coronavirus 229E (CoV-229E). Moreover, we demonstrate that our method can be used in phenotypic drug screening using a panel of nine host- and virus-targeting antivirals. Treatment with effective antiviral compounds reversed the morphological profile of the host cells towards a non-infected state.

**Conclusions:** The method can be used in drug discovery for morphological profiling of novel antiviral compounds on both infected and non-infected cells.

## Background

In the light of the current Coronavirus disease 2019 (COVID-19) pandemic caused by severe acute respiratory syndrome coronavirus 2 (SARS-CoV-2), the development of new therapies against emerging viruses is of high priority. Understanding how viruses affect the host cells is key to identify potential targets for treatment. RNA viruses are characterized by their fast mutation rate resulting in resilience against antiviral drugs, as well as heavy reliance on host pathways for replication (1,2). In this context, coronaviruses are an exception to this norm given the existing RNA proofreading machinery within their genome, but nevertheless develop treatment escape mutants (3). Therefore, host targeting- as well as combination- therapies might be advantageous to battle current and future virus outbreaks. Hence, new methods to profile cellular responses during virus infection that can be used for drug screening are greatly needed.

Image-based morphological profiling of cells, or phenomics, combines high-content imaging with multiparametric analysis of single-cells to study biological and chemical perturbations (4). The Cell Painting assay is a high-content image-based method for morphological profiling that uses six multiplexed fluorescent dyes imaged in five channels. These dyes reveal eight relevant cellular components that can be used to simultaneously interrogate numerous biological pathways upon a given perturbation (5). The resulting images can be analyzed using traditional image analysis techniques to provide information-rich profiles of individual cells, as well as serve as input for machine or deep learning approaches (6). This method has been successfully applied to study compound toxicity, predict cell health indicators, detect morphological disease signatures, and give insights into the mechanism of action (MoA) of both existing and novel compounds (7–12).

In contrast to target-based screens, where only a limited number of features are quantified to select for a known cellular phenotype, morphological profiling combined with statistical analysis can unravel subtle morphological patterns and provide insights into new pathways or mechanisms (5). While morphological profiling is gaining ground in the field of pharmacology and toxicology, its employment in the field of virology has been limited (13). However, with the increased momentum in virology research caused by the spread of the SARS-CoV-2, morphological profiling is progressively being implemented as a strategy for the selection and optimization of drugs, and drug combinations, as potential antiviral therapies (14–16).

Immunofluorescence-based antiviral screening methods commonly rely on the use of specific antibodies for the visualisation and quantification of virus-infected cells. These methods have been proven successful for identifying drugs that directly affect cell survival and infectability of the virus, but lack the depth to provide detailed information about the effect on the host cell (17,18). Here we describe a novel phenomics approach, combining morphological profiling with antibody-based detection of virus infection in a single assay using Human coronavirus 229E (CoV-229E) infected MRC-5 primary lung fibroblasts, in the absence or presence of several known and novel antiviral compounds. We developed an automated image analysis pipeline for the identification of infected and non-infected cells, generating hundreds of morphological measurements to study host-cell biology at a single-cell level. We used the resulting morphological profiles to capture a virus-induced phenotypic signature, we show how these profiles can be used to screen for antivirals that reverse the cellular phenotype from an “infected”- towards a “non-infected” status, and to potentially identify the MoA of novel antiviral compounds.

## Results

### Cell Painting features capture effects of viral infection

In order to investigate how viral infection affects the host cells, the original morphological profiling protocol known as Cell Painting (5) was modified and combined with antibody-based staining. Incorporation of an antibody against virus nucleoprotein (NP) allowed for the identification of virus infection at a single-cell level. In parallel, hundreds of parameters were measured from the infected cells using five of the Cell Painting dyes. Specifically, the dyes consisted of Hoechst (DNA), SYTO 14 (nucleoli and cytoplasmic RNA), Phalloidin (F-actin), Concanavalin A (mannose residues of glycoproteins, especially Endoplasmic Reticulum (ER)), and Wheat Germ Agglutinin (sialic acid and N-acetylglucosamine moieties of glycoproteins, especially plasma membrane and Golgi) (**Fig. 1A and Table 1**). Upon CoV-229E infection, MRC-5 cells were stained using our modified Cell Painting assay combined with a coronavirus NP-specific antibody.

**Figure 1.**
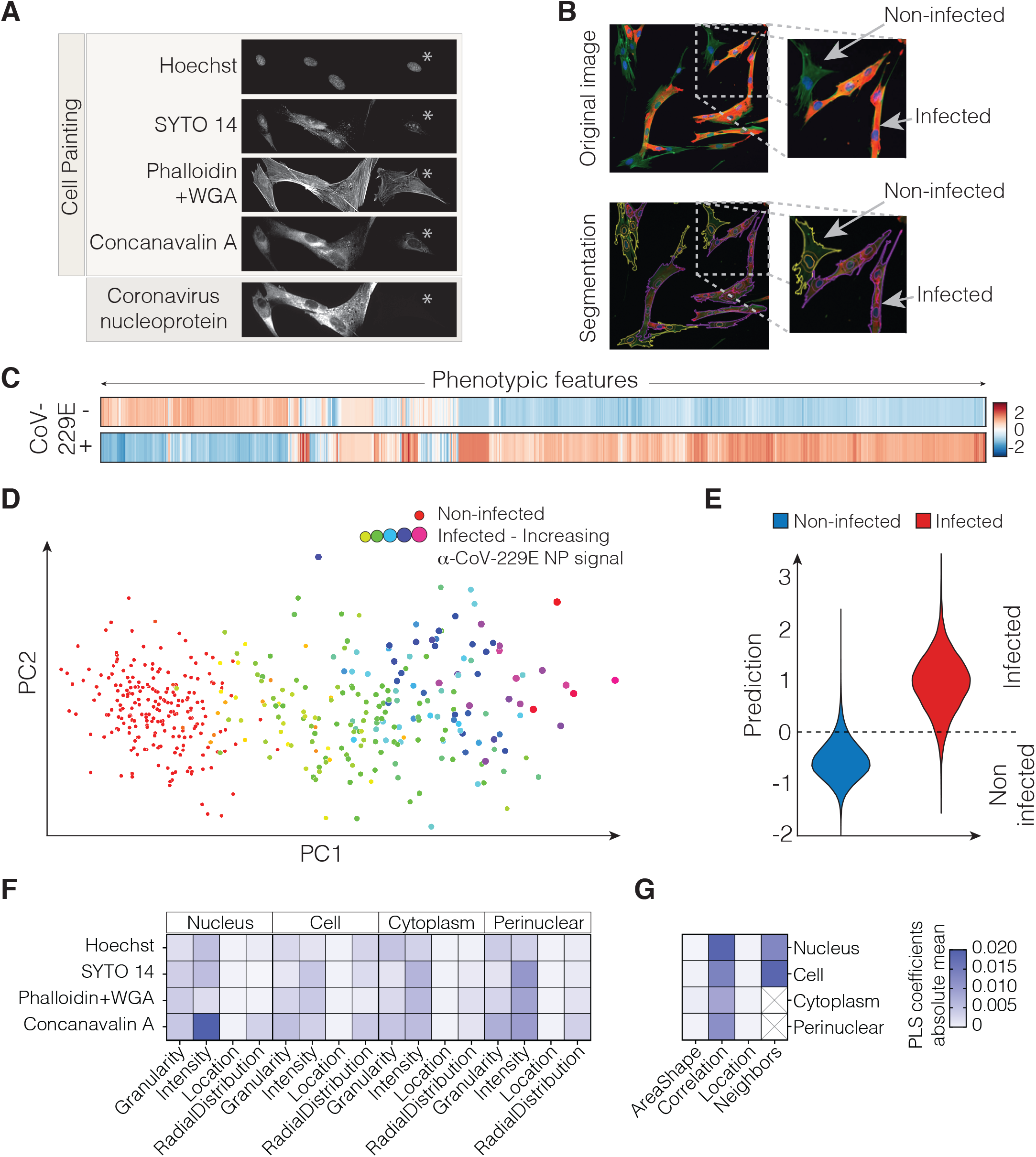
A modified Cell Painting protocol captures a virus-specific morphological signature. **A.** MRC-5 lung fibroblast cells infected with Human coronavirus 229E, stained using Hoechst, SYTO 14, Concanavalin A, Wheat Germ Agglutinin, and Phalloidin, in combination with an anti-coronavirus nucleoprotein (NP) antibody. Note the presence of non-infected (asterisk) and infected cells. **B.** A representative composite image of infected cells with F-actin in green, nuclei in blue and anti-coronavirus NP antibody in red. Segmentation and classification of individual cells visualized with an outline with infected cells in purple and non-infected cells in yellow. **C.** Morphological profiles of non-infected and infected cells (corresponding to the median profiles of both classes). **D.** Dimensionality reduction using PCA applied to the extracted Cell Profiler features per image, coloured according to their infected or non-infected classification based on NP-specific antibody staining. **E.** The PLS-DA prediction model could predict viral infection on cell painting features with substantial accuracy as illustrated by the plot for observed vs predicted values, where observed values correspond to classification by NP-specific antibody. **F,G.** Absolute means of PLS-DA loadings indicate the importance of different feature classes associated with viral infection. Features are grouped according to module, cell compartment, and stain if applicable.

**Table 1.** Problem-solving strategies that separate qualitative/pictorial steps from mathematical steps.

| Label | Excitation spectra | Emission spectra | Provider | Cat-No | Stock concentration | Concentration used | Target organelle |
| --- | --- | --- | --- | --- | --- | --- | --- |
| Hoechst 33342 | 377/50 | 447/60 | Invitrogen | H3570 | 10 mg/ml | 10 µg/ml | nucleus |
| Coronavirus pan Monoclonal Antibody (FIPV3-70) | - | - | Invitrogen | 11500123 | 1 mg/mL | 0.736111111 | Coronavirus viral nucleoprotein |
| Goat Anti-Mouse IgG H&L secondary antibody | 628/40 | 692/40 | Invitrogen | 10739374 | 2 mg/mL | 0.388888889 |  |
| Wheat Germ Agglutinin | 562/40 | 624/40 | Invitrogen | W32464 | 5 mg/m in PBS | 15 µg/ml | Golgi, plasma membrane, |
| Phalloidin | invitrogen | 10135092 | 40x (200U) | 10 µl/ml | F-actin |  |  |
| SYTO 14 green | 531/40 | 593/40 | Invitrogen | S7576 | 5mM in DMSO | 4 µM | nucleoli, cytoplasmic RNA |
| Concanavalin A/Alexa Fluor 488 | 482/35 | 536/35 | Invitrogen | C11252 | 5 mg/ml in 0.1M sodium bicarbonate | 80 µg/ml | Endoplasmic reticulum |

A CellProfiler image analysis pipeline was designed to distinguish between infected and non-infected cells using the coronavirus NP-specific antibody signal in the perinuclear space as a measure (**Fig. 1B**). Using this pipeline, a total of 1441 features corresponding to the five fluorescent dyes were extracted at single cell level, subsequently the cells were labelled according to their infection state based on the NP antibody. After pre-processing, the centred and normalized features were visualized using a heatmap. Clustering of the features highlighted the distinct morphological signature between non-infected and infected cells (**Fig. 1C**). Principal Component Analysis (PCA) was used to reduce redundancy and correlation of the features and to facilitate interpretation. PCA analysis revealed a clear separation between infected and non-infected cells on the first principal component, based on the Cell Painting dyes only (**Fig. 1D**). A gradually increasing separation between non-infected and infected cells coincided with an equally increasing NP antibody signal **Fig. 1D**).

To explore what features contributed most to the virus-induced phenotype, we applied Partial Least-Squares Discriminant Analysis (PLS-DA) using the antibody signal as a label. The PLS-DA model could predict viral infection with substantial accuracy (R2 = 0.73) (**Fig. 1E**). The PLS-DA loadings were used to indicate important feature classes associated with the virus-induced phenotype. The virus-induced morphology was characterized by changes in a wide variety of features, distributed over various cell compartments and stains. The most prominent feature groups related to viral infection, corresponded to the Concanavalin A and SYTO14 staining (An overview of the importance of each of the feature classes, grouped by module, cell compartment, and stain if applicable is provided in **Fig. 1F and G**, and **Suppl. Table 1**). Overall, our analysis demonstrated the strength of phenomics profiling to identify morphological changes exclusive of virus-infected cells.

### A novel phenomics approach for the identification of antiviral compounds

To assess if Cell Painting combined with NP-antibody staining could be leveraged as a tool to profile or screen for antiviral drugs, we designed a phenomics approach including viral infection, compound treatment, cytochemistry protocol, image analysis pipeline, and finally data analysis and visualization (**Fig. 2**, **Suppl. Fig. 1**). This new method is designed to capture in-depth morphological profiles of the host cells induced by virus infection as well as treatment with potential antiviral compounds.

**Figure 2.**
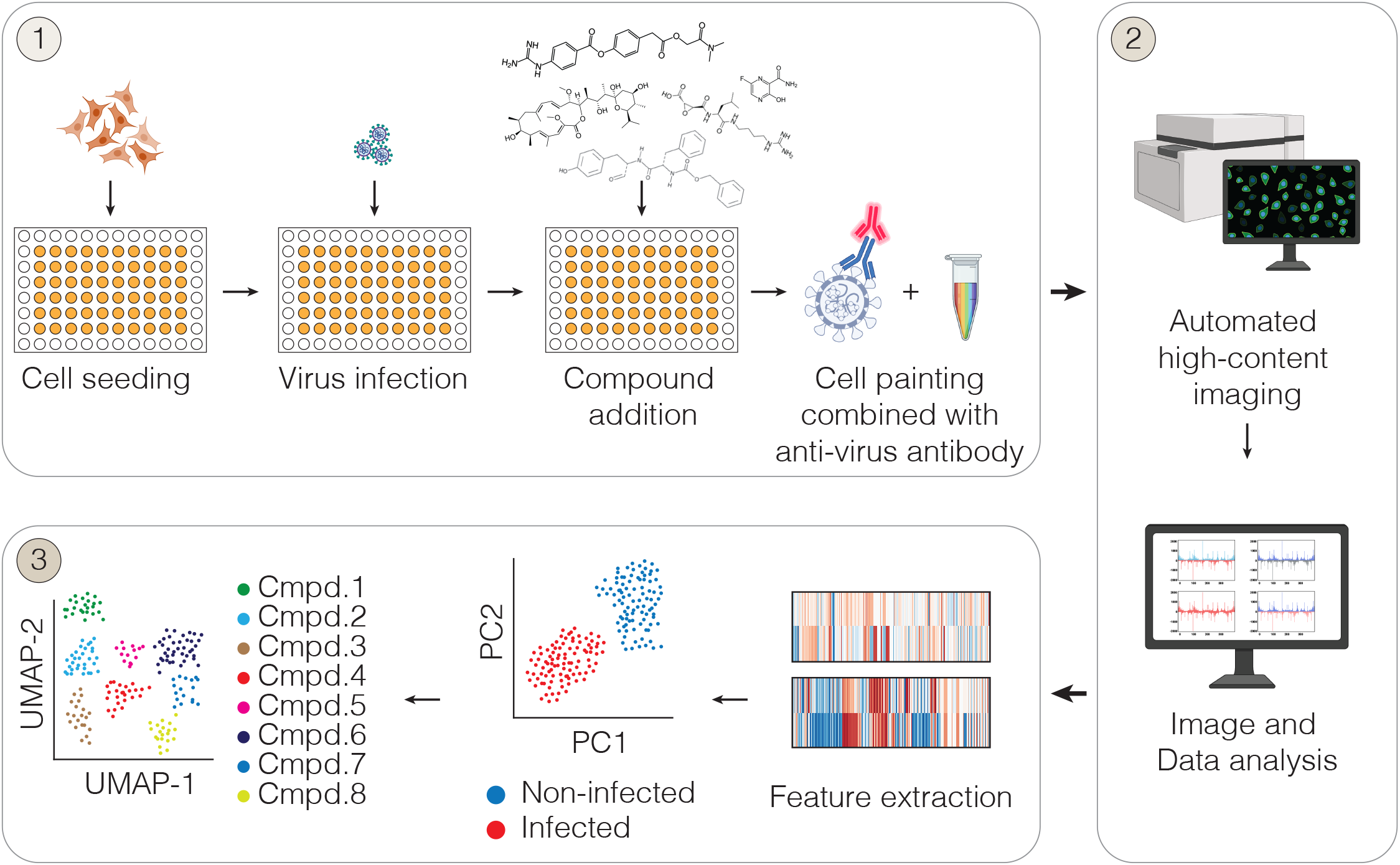
Morphological profiling of virus-infected cells. Overview of the phenomics approach. Cells are seeded in multiwell plates, incubated overnight followed by virus infection. Subsequently, virus-containing media is removed and compounds are added and incubated for 24 or 48h followed by fixation. Then, antibody staining and Cell Painting is performed, followed by high content imaging, image analysis and data analysis of the extracted phenotypic features (a more detailed description of the method can be found in **Suppl. Fig. 1**). Illustrations were partially created with BioRender.com.

For the first experimental section (**Fig. 2A-1**), cells are seeded in multiwell plates and allowed to attach overnight, this is followed by virus infection for one hour. Subsequently, virus-containing media is removed and compounds are added and incubated for 24 or 48h followed by fixation. Next, cells are permeabilized and blocked before the primary antibody against virus NP is applied. After incubation with the primary antibody, a cocktail of Cell Painting dyes and the secondary antibody is added to the cells.

For the second section of the method (**Fig. 2A-2**), high-content imaging is used to capture multiple fields of interest covering the wells. A CellProfiler image analysis pipeline is built to extract single-cell features from all the acquired images. The pipeline includes quality control of the images, pre-processing and feature extraction, and is universal, requiring only minor adjustments for accurate segmentation and determination of thresholds when applied on different cell lines and viruses. For the third and final section of the method (**Fig. 2A-3**), a data analysis and data visualisation pipeline is built. Given that data analysis and representation can be subject to own-interpretation, the work presented here doesn’t provide a specific data analysis pipeline but provides an example of how the data can be analyzed and visualized.

**Figure 3.**
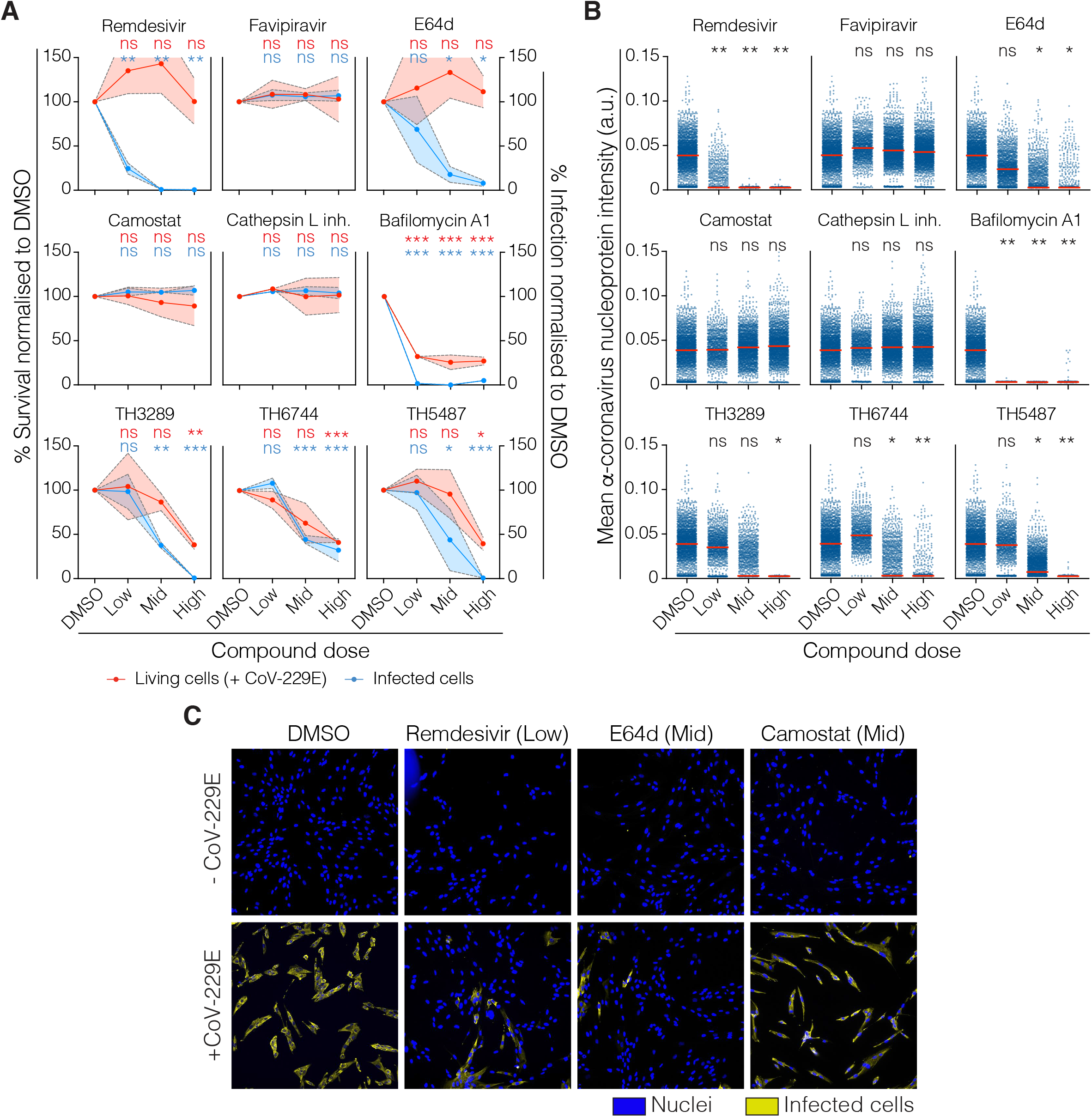
Antiviral efficacy assessment. **A.** Percentage of survival (red) and percentage of infected cells (blue) normalised to DMSO (infected) control. Coloured intervals indicate the standard deviation of two biological replicates. Two-way ANOVA was performed to assess the statistical significance of each condition (*p<0.05, **p<0.005, ***p<0.0005). **B.** Distribution of intensity of the coronavirus anti-NP signal at single-cell level for each tested compound. One-way ANOVA on the means of biological replicates was performed to assess the statistical significance of each condition (*p<0.05, **p<0.005, ***p<0.0005). **C.** Representative images of MRC-5 cells non-infected, or infected with CoV-229E. Hoechst staining for nuclei is visualized in blue, and anti-coronavirus nucleoprotein antibody against CoV-229E in yellow.

### An antibody-based method identifies Remdesivir and E-64d as potent antiviral compounds

To compare our phenomics approach to one of the traditional methods for antiviral drugs screening, we first explored the antiviral efficacy of a set of 9 compounds using an NP-specific antibody. The compounds included two direct-acting drugs targeting viral RNA polymerase (Remdesivir and Favipiravir), and seven host-targeting antiviral compounds (E-64d, Camostat, Cathepsin L inhibitor and Bafilomycin A1), including three novel in-house developed broad-spectrum antiviral compounds TH3289, TH6744 and TH5487 (**Suppl. Fig. 2A**) (19, 20). MRC-5 cells were either not infected or infected with CoV-229E and were then exposed to the compounds at three different concentrations (**Suppl. Fig. 2A**). We determined cell survival by nuclei count and infection rate by the number of anti-coronavirus nucleoprotein positive cells, both in respect to DMSO (**Fig. 3A**). Antiviral activity with no cellular toxicity was observed upon Remdesivir and E-64d treatment. Treatment with clinically utilized Remdesivir reduced infection by 75% at 0.1μM and 99% at 1μM and 8μM concentrations. At the same time, E-64d treatment resulted in a reduction in viral infection of 31%, 82% or 92% compared to DMSO at 1, 10 or 30 μM concentrations, respectively (**Fig. 3A**). Neither FDA-approved Favipiravir nor pre-clinical compounds Camostat or Cathepsin L inhibitor reduced the number of infected cells, while Bafilomycin A1 was toxic at all tested concentrations regardless of the presence of the virus (**Suppl. Fig 2B**). Exposure to the broadly active host-targeting antiviral compounds TH3289, TH6744 and TH5487 showed 62%, 56% and 56% reduction in the levels of infected cells, upon 10 μM treatments, respectively (**Fig. 3A**). The antiviral effect was also confirmed by a decrease in the anti-NP antibody signal (**Fig. 3B and C**), which was evidently lost for the compounds that displayed toxicity.

### Identification of antiviral drugs that reverse infected morphological profiles

In order to demonstrate the use of our approach for drug screening, we assessed the antiviral activity of the aforementioned compounds solely by morphological profiling of the host cell, in the absence of the NP-antibody. We used unsupervised hierarchical clustering to map similarities between the morphological profiles according to their proximity in feature space on all conditions with a survival of 80% or higher. Hierarchical clustering using the Euclidean metric resulted in two main clusters (**Fig. 4A**). The first cluster included both infected cells and infected cells treated with non-effective antivirals (Favipiravir, Camostat and Cathepsin L inhibitor). The second cluster contained both non-infected conditions (control- and compound-treated), as well as infected cells treated with potent antiviral compounds, Remdesivir, E-64d, and the in-house developed TH3289, TH6744 and TH5487, suggesting a rescue or reversion of the virus-induced phenotypic profile (**Fig. 4A**).

**Figure 4.**
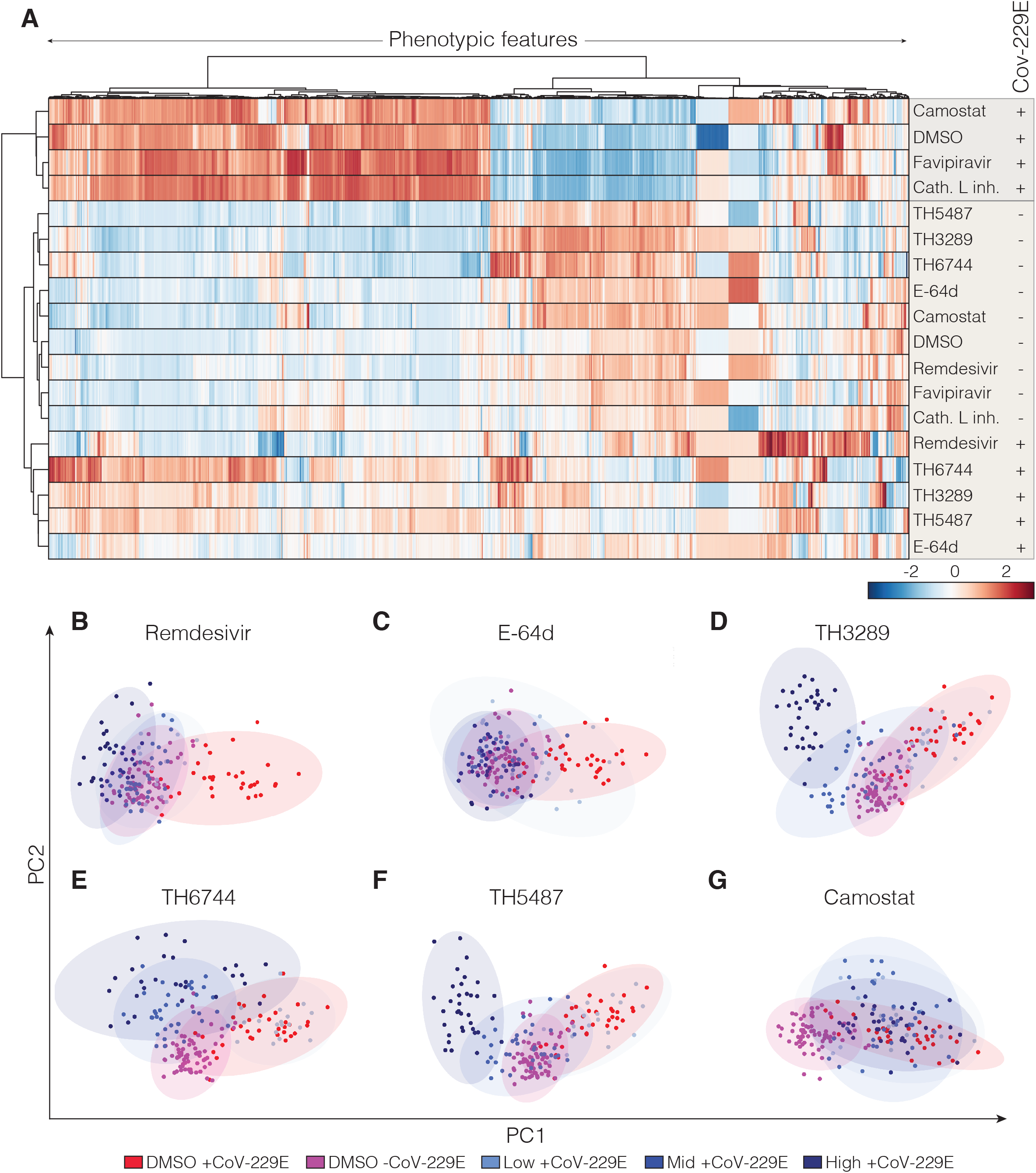
Identification of phenomics profiles of antiviral drugs efficient against coronavirus. **A.** Unsupervised hierarchical clustering of morphological profiles of the indicated samples based on their proximity in feature space. Two main groups are shown, non-infected (-CoV-229E) and infected (+CoV-229E), in the presence or absence of Remdesivir, E-64d, TH6477, Camostat or DMSO (vehicle) at the indicated concentrations. **B to G.** PCA analysis of morphological features upon treatment with Remdesivir, E-64d, TH3289, TH6477, TH5487, and Camostat in non-infected (-CoV-229E) and infected (+CoV-229E) conditions compared to DMSO control. Each dot in the PCA represents one image by taking the mean of all objects in the image.

PCA was used to study the effects of every single compound and dose in more detail. In accordance with the hierarchical clustering, Remdesivir and E-64d treated cells grouped together with non-infected control conditions at all tested doses (**Fig. 4B and C**). Compounds, TH3289, TH6744 and TH5487 showed a dose-dependent response, where treatment with the middle (10 μM) doses showed a trend towards the non-infected cluster, low doses (1 μM) clustered with the infected cells, and high doses (30 μM) formed separate clusters (**Fig. 4D, E and F**). Treatment with the compounds Camostat, Favipiravir and Cathepsin L inhibitor, which did not reduce the number of infected cells, clustered all with the infected cells in PCA space (**Fig. 4G** and **Suppl. Fig. 4A, B**).

The morphological features also provided information on the effect of the compound on the host cells, as illustrated by the morphological profiles in absence of virus (**Suppl. Fig. 3**). The morphological profiles of TH3289, TH6744 and TH5487, which are structural analogues, clustered closely together as indicated by the dendrogram (**Fig. 4A**). Similarly, the TMPRSS family protease inhibitor Camostat clustered together with the protease inhibitor E-64d in the absence of the virus (**Fig. 4A**). Bafilomycin A1 treatment, which was toxic to the cells, as well as cytotoxic doses of the TH compounds, formed clusters separate from both infected and noninfected controls (**Suppl. Fig. 4C**). In summary, these results demonstrate the suitability of our phenomics approach to identify antiviral drugs by reverting virus-induced phenotypic signatures and by providing information on compounds' effect on the host cells.

## Discussion

Traditional targeted screens for antiviral drugs often focus on known proteins in the replication cycle of the virus, neglecting the effects of both the virus and the compound on the host cell. On the contrary, morphological cell profiling enables an untargeted approach for the characterization of cellular responses of infected cells with a high spatiotemporal resolution, high throughput, and on a single-cell level. Here, we present a method utilizing Cell Painting combined with a virus-specific antibody to profile host cell responses upon virus infection and demonstrate its potential to serve as a platform for finding effective antiviral compounds.

By adapting the Cell Painting protocol, we were able to detect a distinct virus-induced phenotype for MRC-5 cells infected with CoV-229E (**Fig. 2C**). The induced phenotype was characterized by changes distributed over various cell compartments and stains, of which the most prominent were observed for Concanavalin A and the SYTO14. The replication cycle of RNA viruses, including Coronaviruses is closely associated with ER and Golgi networks, ensuring close proximity of replication factors, viral protein processing and ultimately viable virus progeny (21). Given that Concanavalin A specifically binds to mannose-rich residues of glycoproteins, which are predominantly present in the ER, the increased intensity of ER signal detected in CoV-229E infected cells can be explained by innate sensing of viral proteins by host ER stress pathways or increased viral glycoprotein content (22). It should be noted that Cell Painting dyes such as Concanavalin A (23) and SYTO-14 are likely to bind to certain viral-induced or viral-derived targets in the host cell, which could in turn contribute to the generated morphological profiles.

Antibody-based screening methods for antivirals have been successfully used in the past to identify potential candidate treatments (24). We used this method to benchmark our phenomics approach. According to their antiviral activity, Remdesivir and E64d, which have been previously shown to ameliorate coronavirus infections (25,26), significantly reduced the percentage of infected cells. Similarly, but with less efficacy, the broad-spectrum antiviral compounds TH3289, TH6744 and TH5487, reduced the number of infected cells, albeit with accompanying toxicity. On the other hand, under our tested conditions, we could not reproduce the previously reported antiviral activities from Favipiravir, Camostat, Cathepsin L inhibitor or Bafilomycin A1 (25–29). These results may be explained by cell-dependent effects, the timing, or the selected doses of treatment, which could be addressed by optimisation of the experimental conditions in future studies.

Despite the fact that antibody-based screening could identify antiviral compounds, it did not provide further information about the host cells beyond viability. Utilizing our phenomics method, we identified antiviral compounds that could reverse the virus-induced morphological profile towards a healthy state. In particular, treatment with Remdesivir, E-64d, TH3289, TH6744 and TH5487 induced morphological profiles that were more similar to non-infected than to infected conditions. Interestingly, as indicated by unsupervised hierarchical clustering, the morphological profiles of TH3289, TH6744 and TH5487, clustered together with drugs that possess much more antiviral efficacy (**Fig. 4A**). Accordingly, this time visualised using PCA, the antiviral activity of Remdesivir and E-64d and the three in-house-developed antivirals was reflected by their clustering close to the control non-infected condition, which indicates a potential rescue of the virus-induced phenotype. Altogether, this points to the fact that based solely on the results of the antibody-based method, the in-house developed compounds would have likely been discarded as antivirals given their toxicity. However, according to the results of our phenomics approach these compounds might have a more potent antiviral efficacy than anticipated, which opens up the possibility for their further development.

A substantial advantage of using morphological profiling to screen for antiviral drugs is the in-depth analysis of the host cell status upon infection as well as upon compound treatment. Thus, despite not being able to recapitulate the antiviral activities of all the tested compounds, we have obtained valuable information that can be used to further understand their biological effect. For instance, in line with several previous studies showing that compounds with similar MoA result in similar phenotypic profiles (4,30,31), for novel compounds such as the ones we tested, the comparison of their morphological profiles against the ones of a reference set could aid at the identification of their MoA. In fact, the unsupervised clustering of the morphological profiles (**Fig. 4A**) seems to indicate that, in the presence of the virus, TH3289, TH5487, and to a lesser extent TH6744, induce a similar signature to E-64d, which despite the need for validation, could suggest that these novel antiviral compounds also have protease inhibitory activity. Despite the very limited number of compounds used in our study, this highlights the use of our method not only to identify opportunities for drugs repurposing but also for the actual discovery of novel antiviral compounds.

A challenge of morphological profiling is the large amount of data generated, which is accentuated when using single-cell measurements. The analysis strategy of such rich data is open to its own interpretation, and might require specific computational and analytical skills. In the case of the work presented here, the statistical and analysis tools used were tailored to our application. However, a comprehensive and detailed guide and toolset on how to analyse morphological profiling data have been previously published (6,11).

Given the increasing implementation of morphological profiling, the biological interpretation of the morphological changes induced by a given perturbation is a subject of active research (4,11). Consequently, the results of this study will benefit from a better understanding of the link between phenomics signatures and the underlying biological mechanism. Interestingly, our results suggest that Cell Painting alone could potentially be used as a phenotypic screening readout to distinguish infected from non-infected cells, and thus be utilized for antiviral screens without the need for an additional virus-specific antibody.

## Conclusions

Our novel phenomics method provides an untargeted readout for the screening of antiviral drugs, in a single assay, at the single-cell level. The morphological signatures obtained using our method can be utilized to identify the potential MoA of novel compounds by comparing them to those of annotated reference drugs, highlighting the use of this method in drug discovery. Morphological profiling provides high adaptability and scalability to study different cell lines and perturbations. Accordingly, we anticipate that our untargeted approach, with only minor adjustments of the image analysis pipeline, will enable other applications using diverse (human-derived) cell lines, as well as different viruses. Overall, we demonstrated that our phenomics approach can be utilized for the unbiased study of virus-induced effects on host cells that can be leveraged for drug repurposing, as well as for the discovery of novel antiviral compounds.

## Methods

### Biosafety

All experiments in presence of infectious virus were performed under respective biosafety laboratory (BSL) conditions according to Swedish Work and Health Authorities. Experiments using CoV-229E were performed at BSL2 at Karolinska Institutet.

### Cell culture

MRC-5 cells (from ATCC, Manassas, VA, USA) were cultured in Minimum Essential Media supplemented with 10% (v/v) fetal bovine serum (FBS; Thermo Fisher Scientific, Waltham, MA, USA), 50 U/ml penicillin and 50 μg/ml streptomycin (Thermo Fisher Scientific, Waltham, MA, USA). The cells were maintained at 37 °C under 5% CO2 and the cell culture was routinely tested for Mycoplasma using a luminescence-based MycoAlert kit (Lonza).

### Virus production

CoV-229E (VR-740; from ATCC, Manassas, VA, USA) stocks were amplified in Huh7 cells. Virus titers were determined in Huh7 cells by end-point dilution assay combined with high-throughput immunofluorescence imaging of viral protein staining as previously described (19).

### Compounds

Remdesivir and Favipiravir were purchased from Carbosynth. TH6744, TH3289 and TH5487 were in-house produced and recently described (19,20). Compounds Camostat (SML0057), E-64d (E8640), Cathepsin L inhibitor (SCP0110), Bafilomycin A (B1793) were purchased from Sigma-Aldrich. Cell painting positive and negative controls included Etoposide (E1383), Fenbendazole (F5396), Metoclopramide (M0763), Sorbitol (S1876) and DMSO (D2438) (Sigma Aldrich).

### Virus infection and compound treatments

Antiviral compounds and cell painting positive controls were dissolved in DMSO to 10mM solutions, and dispersed in 96-well source plates at 500× of the final concentration using the D300e Digital Dispenser (Tecan, Männedorf, Switzerland) and stored at –20 °C until use. Compound and virus infection conditions were randomized within the plates and multiple plate layouts were used to compensate for systematic effects related to the well position. To further avoid plate and edge-effects, the outer wells were excluded from experimentation. Three doses were chosen for each antiviral compound (final concentration; DMSO 0.3% v/v). A total of 2500 MRC-5 cells per well were seeded in black 96 multiwell plates 24 hours before infection (Costar 3603, Corning Life Science, Corning, New York, USA) in 100 μL culture medium. The plates were kept at room temperature for 20 minutes to aid homogeneous spreading in the wells and incubated overnight at 37 °C at 5% CO2 atmosphere. Cells were infected with a MOI 10 or 15 in 50 μL culture medium for one hour and at least four wells per plate were left uninfected as a mock control. Virus-containing medium was removed, replaced with 80 μL culture medium and 20 μL compound-containing medium in at least two technical duplicates, reaching 1× of the final compound concentration. Cells were incubated for 24 or 48 hours. Cells were fixed in 4% paraformaldehyde (Thermo Fisher Scientific, Waltham, MA, USA) in PBS (Gibco, USA) for 20 minutes, washed twice with PBS and stored at 4° C prior to further processing.

### Cell Painting and antibody detection

To profile morphological features of virus-infected cells an adapted version of the cell painting assay was used (5). Specific fluorescent probes or fluorophore-conjugated small molecules were used for the detection of various cellular components (Table 1). To accommodate virus-specific antibody detection in parallel to Cell Painting, the dye specific to mitochondria (MitoTracker) from the original cell Painting protocol was replaced with a coronavirus-nucleoprotein specific antibody. Compared to the original cell painting assay consisting of six dyes targeting eight components/ cell organelles, here five dyes and a virus-specific antibody were used to target virus nucleoprotein in parallel to seven cell organelles and cell components in a single multiplexed staining assay (**Table 1**). Representative images of each of the stains can be found in **Fig 1A.** Coronavirus pan monoclonal antibody (FIPV3-70, Invitrogen) was combined with a secondary antibody using Alexa Fluor 647 fluorophore (10739374, Invitrogen), chosen for its narrow emission and excitation spectra to limit interference with the other channels. To avoid fading of the chemical stains, in our experience, the secondary antibody is best added together with the chemical dyes, but can also be applied sequentially. Concentrations of secondary antibodies need to be adjusted for a good signal-to-noise ratio in the multiplexed assay. In **Table 1**, working concentrations and specifications for the stains and antibodies are presented.

Fixed cells were washed twice (BioTek, 405 LS washer) and permeabilized by incubation with 80 μl of 0.1% Triton X-100 for 20 minutes at room temperature, followed by two washes with 1X PBS. Then, 80 μl of 0.2% BSA (A8022) in PBS was added and incubated for 30 minutes. Next, the plates were washed twice with 1X PBS and 50μl primary antibody was added. The primary antibody was incubated for four hours at room temperature after which the plates were washed three times for five minutes. A staining solution of Hoechst 33342 (Invitrogen, cat.no H3570), SYTO 14 green (Invitrogen, cat.no S7576), Concanavalin A/Alexa Fluor 488 (Invitrogen, cat.no C11252), Wheat Germ Agglutinin/Alexa Fluor 555 (Invitrogen, cat.no W32464) and Phalloidin/Alexa Fluor 568 (Invitrogen, cat.no A12380) was prepared in PBS in addition to Goat anti-Mouse IgG (H+L) Cross-Adsorbed Secondary Antibody, Alexa Fluor 647 (Invitrogen Catalog # A-21235). 50 μl of staining mixture was added to each well and incubated for 30 minutes, after which the plates were washed a final three times for five minutes. Plates were either imaged immediately or sealed and kept at 4 °C to be imaged later. All washing steps were performed using a Biotek 405 LS washer, dispensing was done using the Biotek Multiflo FX, multimode dispenser. Plates were protected from light as much as possible.

### Imaging

Fluorescent images were acquired using an Image Xpress Micro XLS (Molecular Devices) microscope with a 20X objective using laser-based autofocus. In total, 9 sites per well were captured using 5 fluorescence channels (DAPI, Cy5, TexasRed, Cy3 and FITC). To avoid photobleaching, the channels were imaged in order of decreasing excitation wavelength (except DAPI) (**Table 1**). Examples of images from the different channels are shown in **Figure 1A**.

### Image Analysis pipeline

#### Image Quality control and preprocessing

Images were processed and analysed with the open-source image analysis software CellProfiler (available at https://cellprofiler.org/), version 4.0.6 (32). Prior to analysis, a quality control (QC) pipeline was run on the raw images to detect images with artefacts, which may corrupt the data with false values (6). We computed various measures to represent a variety of artefacts and used statistical analysis to detect outliers. Images deviating more than 5 standard deviations from the median of FocusScore, MaxIntensity, MeanIntensity, PercentMaximal, PowerLogLogSlope and StdIntensity were flagged, inspected and removed if necessary. To remove blurred images, the PercentMaximal score and PowerLogLogSlope was computer and images with percentMaximal values higher than 0.25, or PowerLogLogSlope values lower than −2.3 were removed from the dataset. The QC detected 95 images as outliers, as indicated by one or multiple of the control measures. To assess the reproducibility of the assay and detect possible plate and drift effects across multiple plates the various quality measures were visualized and inspected (**Supp. Fig. 5**).

In addition, we used cell-level quality control based on the Hoechst staining to detect bright artefacts and clumped cells. Bright artefacts in the DAPI channel were identified using the Identify Primary Object module in CellProfiler, artefacts and surrounding pixels were masked to avoid interference with segmentation.

To correct for uneven illumination in the images, the polynomial illumination correction function was calculated for each plate and each image channel across all cycles resulting in one illumination correction image per channel. The illumination functions were then used in the analysis pipelines to correct each image by dividing by the respective illumination correction image.

### Segmentation

Segmentation based on the DAPI channel was performed by applying a gaussian blur followed by Otsu thresholding to segment the outlines of each nucleus. The cell objects were segmented using Watershed segmentation using minimum Cross-entropy of the cytoplasmic RNA stain, using the nuclei as a seed. The cytoplasm cell compartment was defined as the cell object subtracted from the nuclei. Segmentation of the perinuclear cell compartment was accomplished by the expansion of 20 pixels in the concanavalin A channel and subtraction of the nuclei. Minor adjustments of the parameters are needed to be used between different experimental setups and on different cell lines.

### Feature extraction

After identification of the four cell compartments, phenotypic characteristics were measured using the AreaShape, Correlation, Intensity, Granularity, Location, Neighbors, and RadialDistribution modules as provided by CellProfiler (32). A total of 1441 features were extracted from each object which were exported into csv format for downstream analysis.

### Classification

To classify the cells for infection state, we used a virus NP-specific antibody. The classification module by CellProfiler Analyst was used to find highly indicative features associated with infection by manual annotation of infected and noninfected cells. Accordingly, the mean intensity of the antibody signal in the perinuclear cell compartment was selected for classification of MRC-5 cells infected with CoV-229E. The threshold for infection was defined as the mean plus three standard deviations from the virus NP-specific antibody intensity in the blank control. The set-off was determined on a plate-to-plate basis to avoid bias by batch-effects and was checked for accuracy by checking for false positives in non-infected wells.

### Antiviral dose-response and cytotoxicity

Nuclei count was used to determine the toxicity induced by compounds alone, as well as compound and virus combined. Averaged nuclei count per condition were normalised to the DMSO controls. To capture relevant cellular phenotypes, cytotoxic compounds resulting in less than 80% survival, were removed from the analysis. Antiviral efficacy was calculated as the percentage of virus-positive cells per well and was normalized to DMSO+virus control which was set to 100%.

### Statistics

All experiments were performed in two separate biological replicates, each consisting of duplicate conditions. Where indicated, plots represent mean +/− SD of the biological replicates. Statistical analysis was done using GraphPad Prism version 9.0 (GraphPad Software Inc). One- or Two-way-ANOVA was performed for all the datasets that required comparison among multiple data points within a given experimental condition. Dunnet was applied for the correction of multiple comparisons, and the family-wise alpha threshold and confidence level were 0.05 (95%confidence interval).

### Feature analysis

In order to facilitate the broad use of our phenomics approach, an image analysis pipeline is provided with the paper, which is tailored to the application here presented, and once executed will provide the necessary input for subsequent analysis. Downstream analysis of the feature data was performed using Python 3. Prior analysis, the data was cleaned from images with artefacts as identified by the QC pipeline described above. Next, the feature values were centred and normalized. Normalization was done across all plates. Invariant features, features with extreme variation (>15 standard deviations) and features with missing data were removed. The features relating to the virus antibody were removed prior analysis of virus-induced morphological profiles.

To compare morphological profiles, the features were presented as cluster maps. Features were scaled using the sklearn StandardScaler method, by subtracting the mean and scaling to unit variance. This ensured that each feature (clustermap column) has a mean of 0 and variance of 1. Unsupervised hierarchical clustering on the morphological profiles was computed using the Euclidean metric, and the clustering algorithm using Ward's method and visualized using the seaborn data visualization library (version 0.11.0).

Visualization of the high dimensional data was done using Principal Component Analysis (PCA) on the centered and normalized features. The image-level feature data was averaged to simplify visualization. Partial Least-Squares Discriminant Analysis (PLS-DA) was used to model the virus infection and select features associated with viral infection. Q2 and R2 values were calculated for the PLS-DA model, which were estimated by five-fold cross-validation. The absolute mean of PLS-DA loadings were calculated and were grouped by Cell Profiler module (AreaShape, Correlation, Intensity, Granularity, Location, Neighbors, RadialDistribution), cell compartment (nuclei, perinuclear space, cytoplasm, cell) and by stain (Hoechst, Concanavalin A, SYTO14, WGA and phalloidin).

## Data availability

All image data, the image analysis pipelines (Quality control, illumination correction and feature extraction), extracted features, as well as an example pipeline are publicly available at FigShare: https://doi.org/10.1101/2021.01.13.423947

## Author contributions

J.R., M.T., A.P., M.-R.P. and J.C-P. designed experiments. M.T. and A.P. performed experiments with BSL2 viruses. J.R and P.G performed morphological profiling experiments, J.R and P.G, M.L performed image analysis, M.L, J.R and J.C-P performed data analysis. O.S. and J.C-P. initiated the project. J.R., M.T., M.-R.P. and J.C-P. wrote the manuscript which was revised by all authors. All authors have read and agreed to the published version of the manuscript.

## Acknowledgements

We thank Professor Tomas Bergström for providing the CoV-229E virus.

## Supplementary information

**Supplementary Figure 1. Detailed overview of the phenomics approach here described.**

**Supplementary Figure 2. Survival of cells exposed to antiviral compounds.**

**A.** List of the drugs and compounds with their corresponding MoA and doses used in this work.

**B.** Nuclei count was used to assess the survival of MRC-5 cells exposed to the indicated compounds for 48h. Two-way ANOVA was performed to assess the statistical significance of each condition (*p<0.02, **p<0.002, ***p<0.0002).

**Supplementary Figure 3. Morphological profiles presented as heatmaps of every tested compound at non-toxic concentrations.**

**Supplementary Figure 4. PCA of the morphological features of each indicated compound.**

**A, B and C.** PCA of morphological features upon treatment with Favipiravir, Cathepsin L inhibitor and Bafilomycin A1 in infected (+CoV-229E) conditions compared to DMSO control (-CoV-229E). Each dot in the PCA represents one image by taking the mean of all objects in the image.

**Supplementary Figure 5. Quality control measures for all plates in the experiment.**

Images deviating more than five standard deviations from the median of FocusScore, MaxIntensity, MeanIntensity, PercentMaximal, PowerLogLogSlope and StdIntensity were flagged as outliers and removed from the analysis.

**Supplementary Table 1. PCA (Fig 1D) loadings of the first Principal component indicating feature importance.**

The top 20 positively correlated, as well as 20 most negatively correlated features including their loadings on the first principal component.

