## Supplementary figures and images for "A phenomics approach for *in vitro* antiviral drug discovery"

### Supplementary Figure 1

Rietdijk J *et al.* Suppl.**Fig.1**

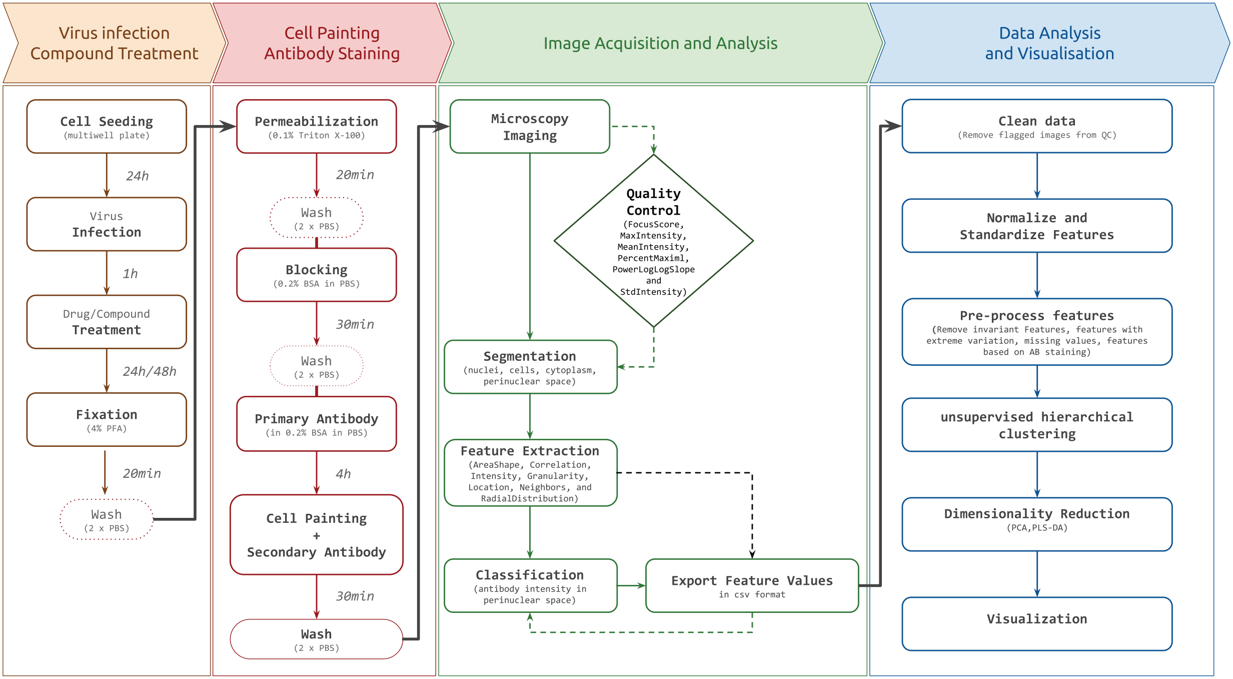

### Supplementary Figure 3

Rietdijk J *et al.* Suppl.**Fig.3**

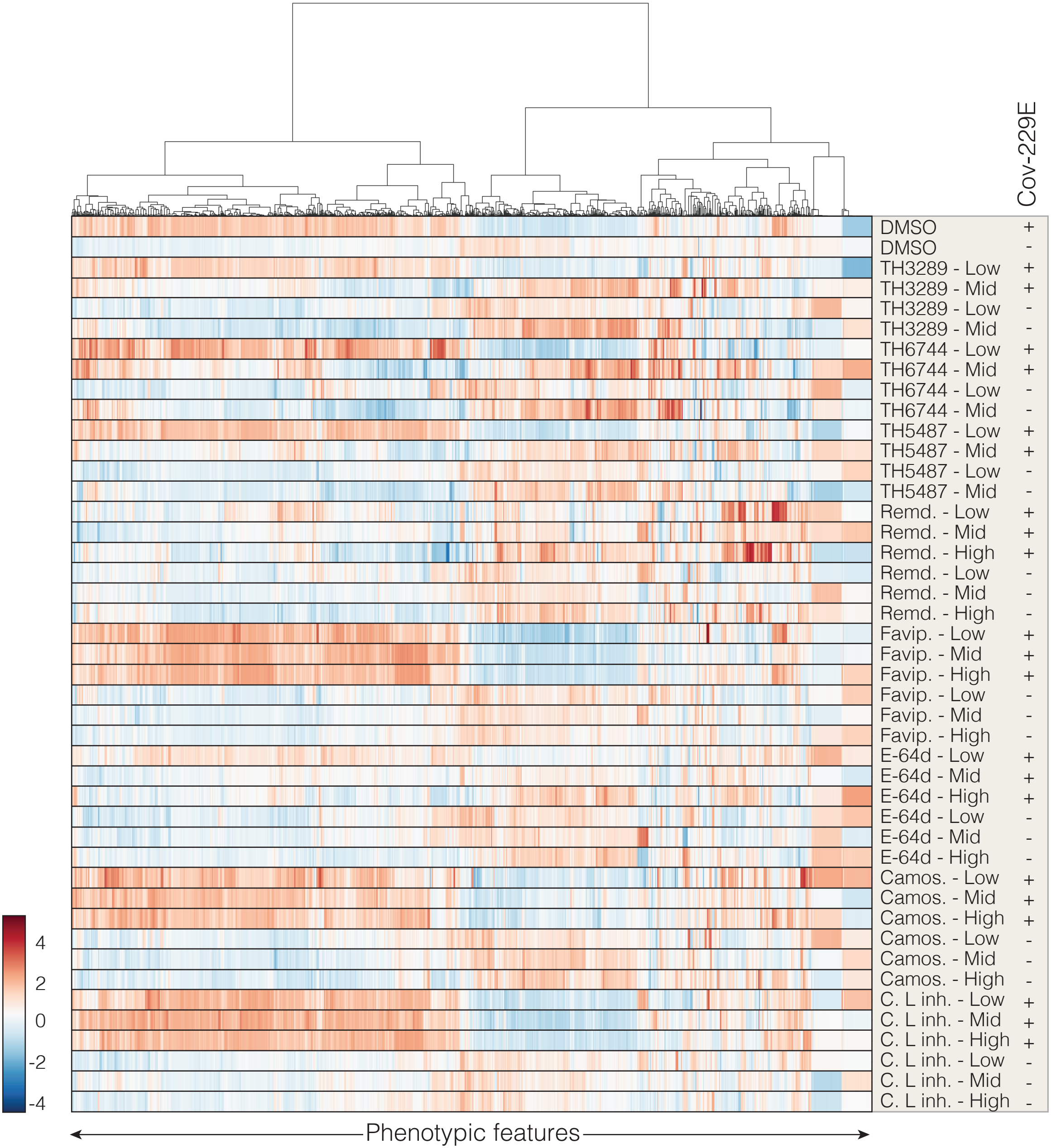

### Supplementary Figure 4

Rietdijk J *et al.* Suppl.**Fig.4**

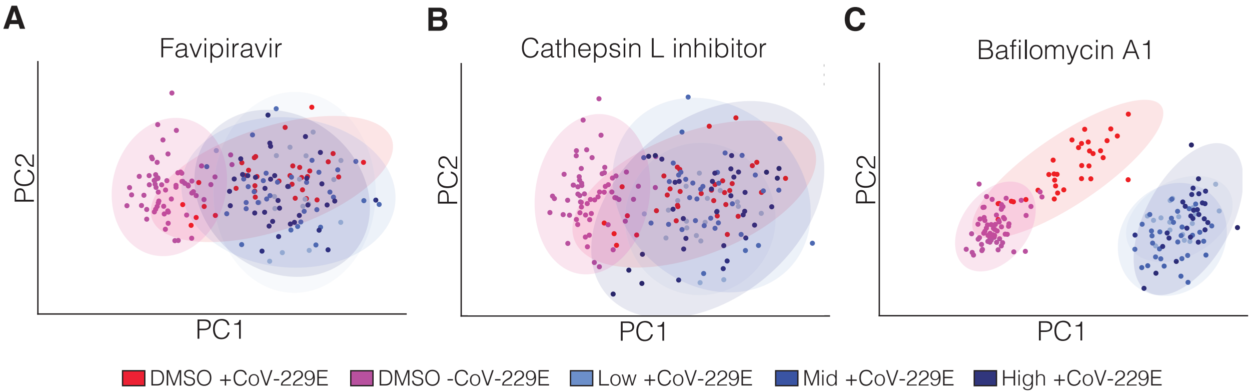

### Supplementary Figure 5

Rietdijk J *et al.* Suppl.**Fig.5**

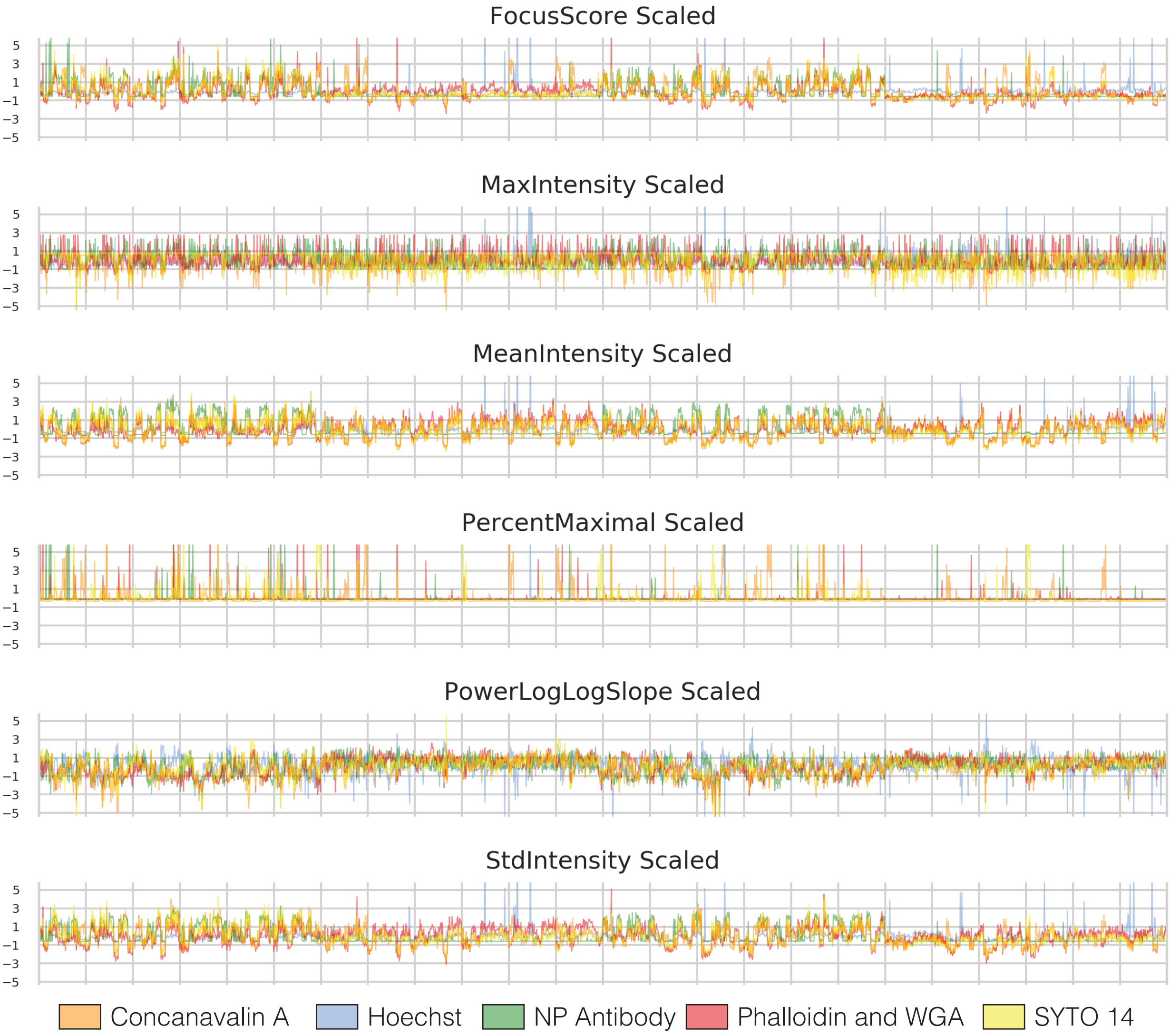
