## Supplementary Figure 2 for "A phenomics approach for *in vitro* antiviral drug discovery"

**A**

| Compound/drug name | MoA | Dose (μM) Low | Mid | High |
|---|---|---|---|---|
| Remdesivir | Nucleoside guanine analogue | 0.1 | 1 | 8 |
| Favipiravir | Nucleotide analogue | 10 | 15 | 30 |
| E-64d | Protease inhibitor | 1 | 10 | 30 |
| Camostat | TMPRSS family protease inhibitor | 0.2 | 10 | 30 |
| Cathepsin L inhibitor | Lysosomal cysteine proteinase | 0.2 | 10 | 30 |
| Bafilomycin A1 | Vacuolar V-ATPase type inhibitor | 0.1 | 1 | 3 |
| TH3289 | Host-targeting antiviral | 1 | 10 | 30 |
| TH6744 | Host-targeting antiviral | 1 | 10 | 30 |
| TH5487 | Host-targeting antiviral | 1 | 10 | 30 |

**B**

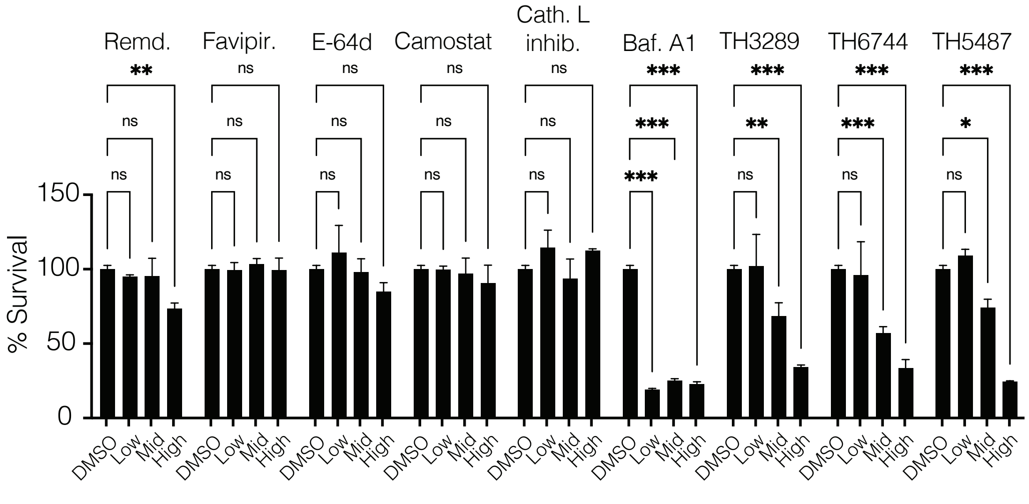
