## Supplementary Table 1 for "A phenomics approach for *in vitro* antiviral drug discovery"

**Supplementary Table 1. Loadings of the first Principal component indicating feature importance.**

| Positively correlated loadings | PC1 | | Negatively correlated loading | PC1 |
|---|---|---|---|---|
| MeanIntensity_Cell_SYTO | 0.074255581 | | AreaShape_EquivalentDiameter_cell | -0.0526401 |
| UpperQuartileIntensity_cell_SYTO | 0.074100557 | | AreaShape_Perimeter_cytoplasm | -0.0523397 |
| MeanIntensityEdge_cytoplasm_SYTO | 0.072543433 | | AreaShape_MaxFeretDiameter_cell | -0.0522439 |
| UpperQuartileIntensity_cytoplasm_SYTO | 0.072466699 | | AreaShape_MaxFeretDiameter_cytoplasm | -0.0522439 |
| MeanIntensity_cytoplasm_SYTO | 0.072217955 | | AreaShape_EquivalentDiameter_cytoplasm | -0.0519756 |
| MedianIntensity_cell_SYTO | 0.071093266 | | RadialDistribution_FracAtD_perinuclear | -0.0516869 |
| StdIntensityEdge_cytoplasm_SYTO | 0.071005105 | | AreaShape_Perimeter_cell | -0.0512184 |
| MeanIntensityEdge_nuclei_SYTO | 0.070791576 | | AreaShape_MajorAxisLength_cytoplasm | -0.0501069 |
| UpperQuartileIntensity_perinuclear_SYTO | 0.070453333 | | AreaShape_MajorAxisLength_cell | -0.0499238 |
| MeanIntensity_perinuclear_SYTO | 0.070204563 | | RadialDistribution_FracAtD_perinuclear | -0.0492749 |
| MeanIntensityEdge_perinuclear_SYTO | 0.069817079 | | RadialDistribution_FracAtD_perinuclear | -0.0486969 |
| MedianIntensity_perinuclear_SYTO | 0.068761953 | | RadialDistribution_FracAtD_perinuclear | -0.0476608 |
| MeanIntensityEdge_cytoplasm_CONC | 0.068554908 | | AreaShape_CentralMoment_cell | -0.046758 |
| MedianIntensity_cytoplasm_SYTO | 0.068303186 | | AreaShape_SpatialMoment_cell | -0.046758 |
| MedianIntensity_nuclei_CONC | 0.067808394 | | AreaShape_Area_cell | -0.046758 |
| MeanIntensity_nuclei_CONC | 0.067732508 | | RadialDistribution_MeanFrac_cytoplasm | -0.0467017 |
| UpperQuartileIntensity_cell_PHAandWGA | 0.067605708 | | Granularity_1_cytoplasm | -0.0460395 |
| LowerQuartileIntensity_nuclei_CONC | 0.067574965 | | AreaShape_CentralMoment_cytoplasm | -0.0457216 |
| StdIntensity_cytoplasm_SYTO | 0.06746692 | | AreaShape_SpatialMoment_cytoplasm | -0.0457216 |
| LowerQuartileIntensity_nuclei_SYTO | 0.067416544 | | AreaShape_Area_cytoplasm | -0.0457216 |
